# The Neural Correlates of Arousal: The Ventral Posterolateral Nucleus-Global Transient Co-Activation

**DOI:** 10.1101/2022.11.23.517776

**Authors:** Junrong Han, Qiuyou Xie, Xuehai Wu, Zirui Huang, Sean Tanabe, Stuart Fogel, Anthony G. Hudetz, Hang Wu, Georg Northoff, Ying Mao, Sheng He, Pengmin Qin

## Abstract

Arousal and awareness are two components of consciousness whose the neural mechanisms remain unclear. Spontaneous increases of global (brain-wide) blood-oxygenation-level-dependent (BOLD) signal has been found to be sensitive to changes in arousal. By contrasting BOLD datasets with altered arousal levels, we found that the activation of ventral posterolateral nucleus (VPL) decreased during transient increase in the global signal (top 17% data) in low arousal and awareness states (non-rapid eye movement sleep and anesthesia) as compared to wakefulness, and even in eye-closed (compared with eyes-open) in healthy awake-states, while this activation remained unchanged in patients with unresponsive wakefulness syndrome characterized by high arousal without awareness. These results demonstrate that co-activation of the VPL and global activity is critical to arousal, but not to awareness.

**One-Sentence Summary:** The VPL nucleus-global brain transient co-activation is related to physiological arousal but not to perceptual awareness.

## Introduction

Consciousness can been divided along two axes: awareness (i.e. subjective experience; the mental contents of consciousness) and arousal (i.e. the states of wakefulness; neurophysiological arousal) (*1*). In researching these two components, scientists have found, that both slow-wave sleep and anesthesia states have reduced awareness and arousal, while unresponsive wakefulness syndrome (UWS), formerly named as the vegetative state, is characterized by high arousal and low awareness (Fig.1A). Specifically, a high arousal was marked by eye opening (*1*) and higher neural activity of the brain stem (*2*). However, despite an extensive research into awareness (*3–7*), there remains a controversy on how specific arousal systems and large-scale brain activity interact in regulating the level of arousal as a component of consciousness.

**Fig. 1.**
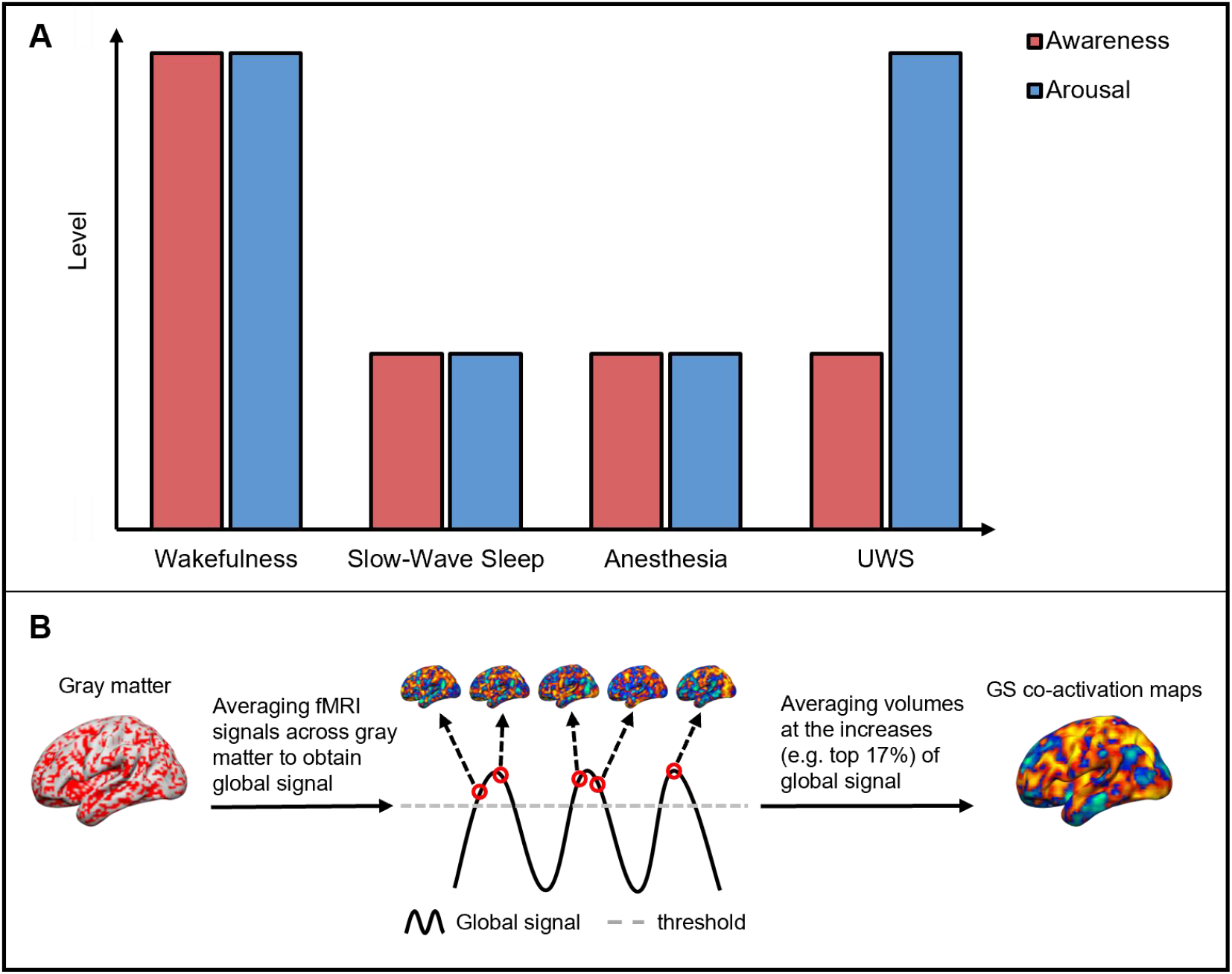
Various arousal states and main analysis we used in this study. (**A**) The simplified illustration of two consciousness components (awareness and arousal) in various states, including wakefulness, slow-wave sleep, anesthesia and UWS patients. (**B**) The schematic of calculating the GS co-activation maps.

Nevertheless, recent evidence suggests that the brain-wide (e.g., global) spontaneous blood oxygenation level dependent (BOLD) signal (GS) fluctuations may be related to arousal (*8–11*). For instance, one study showed that the GS amplitude, defined as the temporal standard deviation of GS time series, showed a cumulative decrease from morning to afternoon, which suggested a positive correlation between arousal and the GS amplitude(*12*) (but see other studies (*8, 9*)). Most interestingly, based on former findings, global spontaneous activity could support arousal in collaboration with the thalamus. For example, the transient increase in GS (top 17% of GS series) during the resting state was found to be associated with the deactivation in the dorsal midline thalamus, which was thought to be related to a momentary drop of arousal (*13*). On the other hand, opposite findings have also been found in that deep brain stimulation of the central thalamus or the central lateral thalamus induced widespread cortical activity, which was accompanied by an improvement of consciousness in heavily sedated macaques and patients with disorders of consciousness (*14–16*). Given that these studies showed that the dorsal midline thalamus, central thalamus, and central lateral thalamus modulated consciousness, which includes both arousal and awareness, it is unlikely that these thalamic nuclei should be exclusively related to arousal but not awareness. This has led to our hypothesis that the neural mechanisms underlying arousal could involve other thalamic nuclei, such as the sensory thalamic nuclei, that modulate global spontaneous activity and have a potential relationship with perceptual consciousness (*17–20*).

To investigate the role of functional interaction between thalamic and global brain activities in arousal, we here analyzed multi-center resting-state fMRI datasets at different arousal levels, including various sleep states, anesthesia, UWS, and wakeful resting-state in healthy participants with eyes open and eyes closed (EO/EC) (Table 1). We first explored how the brain (in particular, thalamus) activity during GS increase changed among strongly altered arousal levels (N2, N3 sleep and anesthesia), and slightly altered arousal levels between eyes-open and eyes-closed conditions in healthy awake individuals. Given that arousal and awareness are reduced simultaneously in normally suppressed states (i.e., sleep and anesthesia), to dissociate these components of consciousness (i.e. awareness and arousal), we analyzed additional data from UWS patients characterized by a high arousal without obvious signs of awareness (*1*).

**Table 1.**
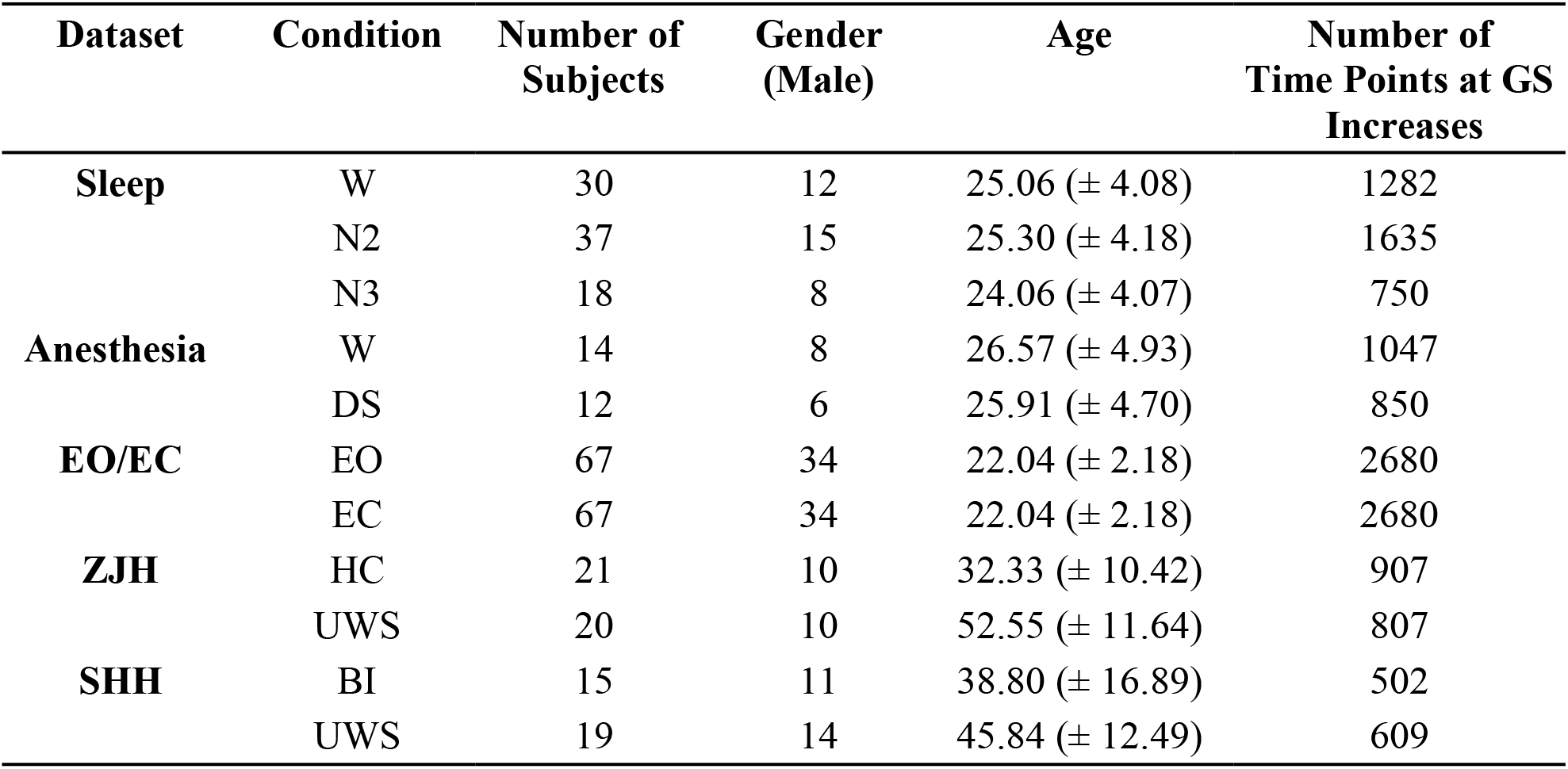
Detailed Information for All Datasets.

Specifically, for all the datasets, we conducted co-activation analysis to capture momentary changes of activity in the thalamus and cortical regions during spontaneous transient increases of the GS (*21*). We first calculated the GS by averaging the z-score transformed from time series of all voxels within the gray matter, and then selected the top 17% of time points of the GS to represent the instantaneous increases (*13, 22*). To minimize the whole-brain effect of various arousal states, the z-score map of each brain volume was normalized by subtracting the GS value and then dividing the result by the GS value. Finally, the normalized activation maps at the time points of the GS increase (top 17%) were averaged to produce the GS co-activation map (*13, 22*) (Fig.1B).

As we will show, increases in GS were accompanied by higher activity in ventral posterolateral nucleus (VPL), a sensory nucleus of thalamus, only in high arousal states irrespective of the level of awareness suggesting that their co-activation is critical to arousal, but not to awareness.

## Result

To investigate how the GS co-activation pattern changes during naturally reduced arousal, we adopted the fMRI data of natural sleep stages which included wakefulness, N2 and N3 stages (Table 1). The GS co-activation maps were compared across three stages using ANOVA. As shown in the figure below, the analyses first showed that the thalamus, auditory cortex (AUD), visual cortex (VIS), and sensorimotor cortex (SMT) were significantly modulated across sleep stages (voxel-wise p < 0.001, cluster-wise alpha < 0.005) (Fig.2A). Region of interest (ROI)-based analysis was further used to compare the GS co-activation on the above regions between each paired stages and found a significantly higher GS co-activation in the thalamus, AUD, VIS, and SMT during wakefulness than N2- and N3-sleep (unpaired t-test, p < 0.001, Bonferroni corrected) (Fig.2A, Fig.S1A). Moreover, to test whether the ROIs showed a distinctive activation pattern at GS increases, their normalized activation at random time points were also examined. The result showed that the normalized activation of these ROIs did not significantly differ between stages at random time points (Fig.2A, Fig.S1B). And the physiological noise, including white matter (WM) and cerebral spinal fluid (CSF), showed no significant difference of GS co-activation among sleep stages (Fig.S1C), which ruled out the possibility that the GS co-activation change in thalamus and other regions were contributed by these noises.

**Fig. 2.**
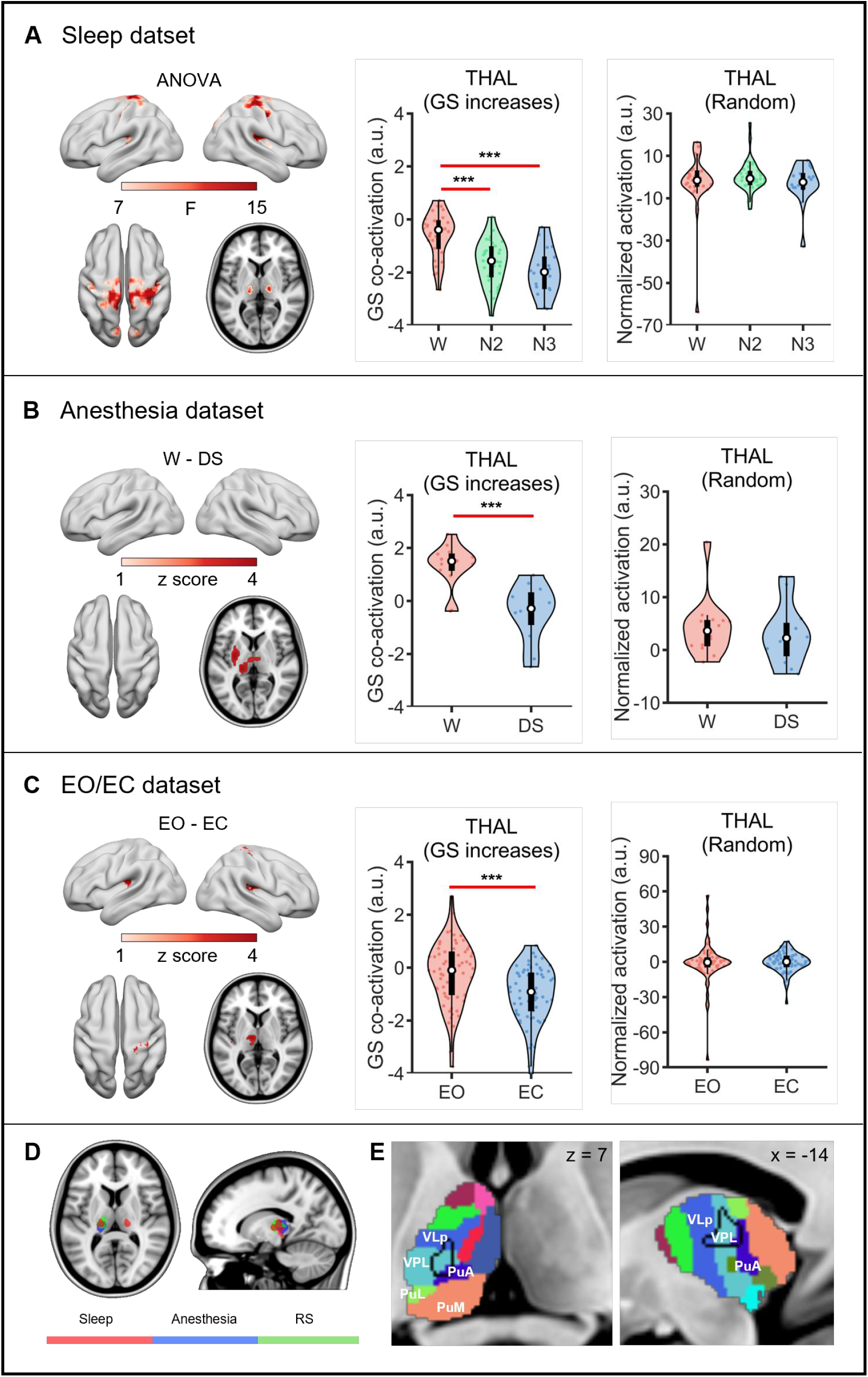
The GS co-activation of VPL decreases in all reduced arousal states. (**A**) Sleep dataset: comparison of GS co-activation maps between wakefulness (W), N2 and N3 sleep (voxel-wise p < 0.001, cluster-wise alpha < 0.005) (left). GS co-activation comparison between stages at the thalamus (THAL) (*** = p < 0.001, Bonferroni corrected) (middle). Comparison of normalized activation during random time points between stages at the thalamus (right). (**B**) Anesthesia dataset: comparison of GS co-activation map between wakefulness (W) and deep sedation (DS) (voxel-wise p < 0.001, cluster-wise alpha < 0.005) (left). GS co-activation comparison between W and DS at the thalamus (middle). Comparison of normalized activation during random time points at the thalamus (right). (**C**) EO/EC dataset: comparison of GS co-activation map between eyes-open (EO) and eyes-closed (EC) (voxel-wise p < 0.001, cluster-wise alpha < 0.005) (left). GS co-activation comparison between EO and EC at the thalamus (middle). Comparison of normalized activation during random time points at the thalamus (right). (**D**) The overlapped region in the thalamus identified by overlapping the resulting brain maps from the above three datasets. (**E**) The overlapped regions in the thalamus further identified as the VPL nucleus using the atlas of subcortical area. Black line indicates the edge of the overlapped thalamus regions in (D).

To further examine whether the thalamus-global transient co-activation was also affected by pharmacological arousal reduction, we adopted an anesthesia dataset which included the wakefulness and deep sedation conditions (Table 1). The unpaired t-test showed significantly higher co-activation in the thalamus and putamen in wakefulness than deep sedation (voxel-wise p < 0.001, cluster-wise alpha < 0.005) (Fig.2B, Fig.S2A). And the normalized activation of these ROIs did not show significant difference between conditions at random time points (Fig.2B, Fig.S2B), neither did the GS co-activation of physiology noise (Fig.S2C).

Furthermore, to examine whether this thalamus-global transient co-activation could be influenced even in slightly reduced arousal, we adopted the resting-state fMRI datasets from healthy participants during eyes-open and eyes-closed conditions (Table 1). The comparison showed significantly decreased GS co-activation in thalamus, AUD and SMT in the eyes-closed condition compared to eyes-open (paired t-test, voxel-wise p < 0.001, cluster-wise alpha < 0.005) (Fig.2C, Fig.S3A), while no significant difference of normalized activation during random time points were found in these regions (Fig.2C, Fig.S3B). Additionally, no significant difference of GS co-activation between conditions was found in physiological noises (Fig.S3C).

Subsequently, by overlapping the brain mappings from anesthesia, sleep and EO/EC groups, only the thalamus was identified as the brain region that instantly co-activated with the GS increase (i.e. the thalamus-global transient co-activation) in the awake state, of which the co-activation was disrupted when arousal decreased (Fig.2D). Further analysis using the atlas of subcortical areas (*23*) pinpointed this thalamus region to be the VPL nucleus (Fig.2E).

To dissociate the different involvement of the thalamus-global transient co-activation in arousal and awareness, we recruited two datasets obtained from Zhujiang Hospital (ZJH) and Shanghai Hospital (SHH), which included UWS patients who were considered to have high arousal without awareness (*1*). To minimize the effects of structural distortion on the subsequent analysis, we only included the UWS patients with well-preserved brain structures(*7*). The ZJH and SHH datasets also contained healthy subjects and fully conscious patients with a brain injury (BI) history as control groups (Table 1). According to our hypothesis, if the GS co-activation of the thalamus was as preserved in the UWS (high arousal without awareness) as in healthy control (HC) or BI, it could be inferred that the thalamus-global transient co-activation is exclusively related to arousal rather than awareness. To test this hypothesis, we averaged the GS co-activation within the overlapped thalamus ROI obtained from previous experiments (Fig.2D) and compared it between UWS and HC (ZJH dataset), and between UWS and BI (SHH dataset). The results showed no significant GS co-activation difference at the overlapped thalamus between UWS and HC (p = 0.14, Bonferroni corrected) (Fig.3A) or between UWS and BI (p = 0.99, Bonferroni corrected) (Fig.3B). Furthermore, the normalized activation of the overlapped thalamus at random time points did not show significant differences between conditions in the two datasets (Fig.3A, B), neither did the GS co-activation of physiological noises (Fig.S4A, B). These results suggested that the transient co-activation between the thalamus (especially, the VPL nucleus) and global cortical activity was less affected in the UWS patients, thus supported our hypothesis that this co-activation should be more related to arousal, than to awareness.

**Fig. 3.**
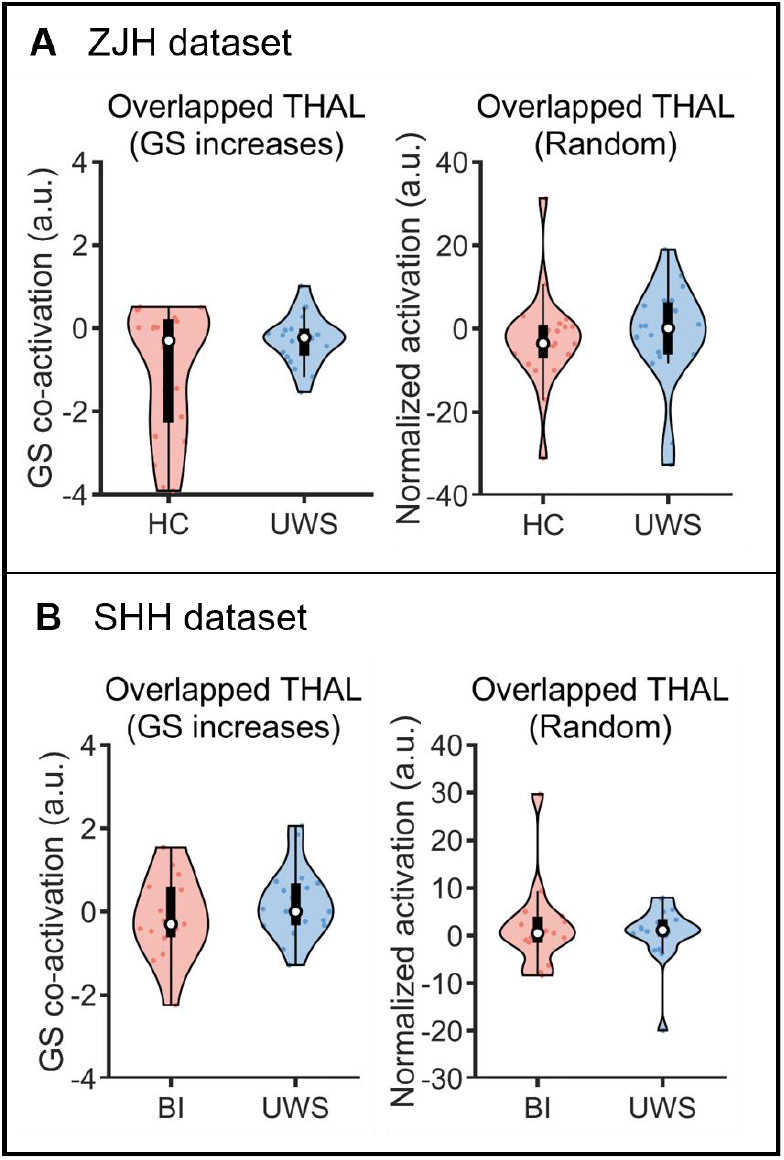
The GS co-activation of VPL remains unchanged in UWS patients. (**A**) ZJH dataset: comparison of GS co-activation at the overlapped thalamus between HC and UWS (left). Comparison of normalized activation at the overlapped thalamus during random time points (right). (**B**) SHH dataset: Comparison of GS co-activation at the overlapped thalamus between UWS and brain injury (BI) (left). Comparison of normalized activation at the overlapped thalamus during random time points (right).

In addition, we performed an ROI-based analysis by averaging the preprocessed BOLD signal during GS increases within the overlapped thalamus and compared it between conditions in each dataset. All the results replicated the above findings, in which the activity of overlapped thalamus (mainly at the VPL nucleus) during GS increases remained high in wakefulness, EO, HC, BI, and UWS, but decreased in N2-sleep, N3-sleep, anesthesia and EC (p < 0.05, Bonferroni corrected) (Fig.S5).

## Discussion

In summary, the current findings showed that the activation in the VPL of the thalamus was higher during the transient GS increase in all the high arousal conditions (e.g. wakefulness, EO, HC, BI, UWS) relative to low arousal conditions (e.g. N2-sleep, N3-sleep, anesthesia, and EC), indicating that the co-activation between the VPL and the whole cortex supported arousal in a general way. Most importantly, the dissociation between arousal and awareness in UWS (high arousal, but low awareness) indicated an exclusive relationship of the high VPL activation during GS increase to arousal, rather than awareness.

The current results showed that the transient GS increase, accompanied with the activity in VPL, supports the arousal levels. This finding is supported by previous studies. For instance, animal studies found that the activity of the sensory thalamic nuclei could modulate the cortical activity levels during quiet wakefulness and non-rapid eye movement sleep (*17–19*). The current results extended the previous findings by showing the role of sensory thalamus nucleus (VPL) in supporting arousal. So, how does VPL, a sensory thalamus nucleus found to receive ascending input from the medial lemniscus and project to the somatosensory cortex (*24, 25*), affect wide-spread cortical activity to modulate the arousal states? One recent study provided clues that the sensory-motor regions could drive the spontaneous large-scale cortical activity switching between a high coherent state and a low coherent state (*20*). Moreover, one recent theory of sleep-wake control proposed an intimate association between arousal system and somatic motor control circuits (*26*). Combined with our results, the VPL could achieve the co-activation with large-scale cortical regions through the sensory-motor regions to implement the arousal modulation. Although both the current results and some previous findings showed that the GS reflected brain-wide (or global) fluctuations in resting BOLD signals and carried neurophysiological information (*22, 27, 28*), some studies proposed otherwise, and regarded GS as a physiological and head-motion noise (*29, 30*). In the current study, we conducted corresponding control analyses to rule out the potential confounding effects from those noises (Fig.S1-3C, Fig.S4).

The current findings showed no difference of VPL activation between the UWS and the fully conscious states (i.e HC and BI), which strongly suggested that the VPL-GS transient co-activation was exclusively related to arousal, but not awareness, as UWS patients are characterized by high arousal without awareness (*1*). In addition to the previous finding of a high activation in the brain stem (*2*) and specific EEG features for arousal in UWS (*31*), our findings showed for the first time, that special connection between subcortical structures and the cortex could be generally related to arousal, especially the arousal state in UWS. More interestingly, this result was also different from the previous animal studies, in which activation of non-sensory thalamic nuclei were found to induce wide-spread cortical activity in anesthesia (or sleeping) monkey and mice, accompanied with the recovery of consciousness (both arousal and awareness) (*14, 15, 32, 33*). In these animal studies, the central lateral thalamus (CL), central thalamus (CT), centromedial thalamus (CMT), and the paraventricular thalamus (PVT) were used as targets in the thalamus, which all receive input from the brainstem reticular activating system, and project to wide-spread cortical regions. Moreover, one human study showed that in patients with disorders of consciousness, stimulation on the CT could also improve the patients’ awareness (*16*). However, all these previous studies focused on the relationship between non-sensory thalamic nuclei and the large-scale cortical regions in supporting consciousness (both arousal and awareness), not arousal alone.

Furthermore, contrary to the positive relationship between thalamic nuclei and the neocotex found in the current study and the above mentioned studies, one recent study showed that the GS transient increase co-occurred with the de-activation in the dorsal midline thalamus and the nucleus basalis, suggesting that the GS increase was related to the momentarily reduced arousal (*13*). This activity is usually found in the microsleep state (*10, 34–36*) or transition from wakefulness to anesthesia (*37*), which was proposed to reflect the cortical activity of the brain’s attempt to prevent the drop of awareness or arousal (*35*). In view of the dynamic contribution of different cortical regions to GS (*38, 39*), and of the diverse relationship between the thalamic nuclei and GS, the global activity, VPL and non-sensory thalamic nuclei are likely to work together in a complex way to support arousal and normal consciousness, which calls for further investigations in the future.

In conclusion, we revealed that the higher activation in the VPL nucleus during transient increase of GS is exclusively related to arousal, which shed lights on our understanding of the neural correlates of arousal and consciousness. Our finding of a possible exclusive influence of VPL on arousal suggested a possibility that electrical stimulation to the sensory nucleus of thalamus could improve the arousal level of anesthetized macaques without improving their awareness, which could provide a new animal model for UWS, to explore novel options in the diagnosis and treatment of disorders of consciousness.

## Acknowledgments

We thank Jinhui Wang and Ning Liu for discussions and critical comments on the manuscript.

## Funding

The Key Realm R&D Program of Guangzhou (202007030005 to PQ)

The National Natural Science Foundation of China (Grant 31971032 to PQ)

The National Natural Science Foundation of China (Grants 82271224 to XW)

Lingang Laboratory (Grant No. LG202105-02-03 to XW)

The Shanghai Science and Technology Development funds (No. 16JC1420100 to Y.M)

The Shanghai Municipal Science and Technology Major Project (No.2018SHZDZX01 to Y.M)

The Natural Science Foundation and Major Basic Research Program of Shanghai (16JC1420100)

Canadian Institutes of Health Research (CIHR)

Michael Smith Foundation (EJLB-CIHR)

The Hope for Depression Research Foundation (HDRF)

The HBP Joint Platform to GN, funded from the European Union’s Horizon 2020 Framework Program for research and Innovation under the specific Grant Agreement No 785907 (Human Brian Project SGA 2)

## Author contributions

Q.X, X.W, Z.H, S.T, S.F collected data, J.H analyzed data, J.H and P.Q designed the study and wrote the manuscript, and all authors edited the manuscript.

## Competing interests

The authors declare that they have no competing interests.

## Data and materials availability

All data that support the findings of this study is available on request from the corresponding author.

## Supplementary Materials

Materials and Methods

Figs. S1 to S5

References (*40–43*)

## Notes

### Competing Interest Statement

The authors have declared no competing interest.

## References

1. S. Laureys, The neural correlate of (un)awareness: Lessons from the vegetative state. Trends Cogn. Sci. 9, 556–559 (2005).

2. J. L. Bernat, Chronic disorders of consciousness. Lancet. 367, 1181–1192 (2006).

3. A. Demertzi, E. Tagliazucchi, S. Dehaene, G. Deco, P. Barttfeld, F. Raimondo, C. Martial, D. Fernández-Espejo, B. Rohaut, H. U. Voss, N. D. Schiff, A. M. Owen, S. Laureys, L. Naccache, J. D. Sitt, Human consciousness is supported by dynamic complex patterns of brain signal coordination. Sci. Adv. 5, 1–12 (2019).

4. Z. Huang, J. Zhang, J. Wu, G. A. Mashour, A. G. Hudetz, Temporal circuit of macroscale dynamic brain activity supports human consciousness. Sci. Adv. 6, 1–15 (2020).

5. A. I. Luppi, M. M. Craig, I. Pappas, P. Finoia, G. B. Williams, J. Allanson, J. D. Pickard, M. Owen, L. Naci, D. K. Menon, E. A. Stamatakis, Consciousness-specific dynamic interactions of brain integration and functional diversity. Nat. Commun. 10, 4616 (2019).

6. A. B. A. Stevner, D. Vidaurre, J. Cabral, K. Rapuano, S. F. V. Nielsen, E. Tagliazucchi, H. Laufs, P. Vuust, G. Deco, M. W. Woolrich, E. Van Someren, M. L. Kringelbach, Discovery of key whole-brain transitions and dynamics during human wakefulness and non-REM sleep. Nat. Commun. 10, 1035 (2019).

7. P. Qin, X. Wu, C. Wu, H. Wu, J. Zhang, Z. Huang, X. Weng, D. Zang, Z. Qi, W. Tang, T. Hiromi, J. Tan, S. Tanabe, S. Fogel, A. G. Hudetz, Y. Yang, E. A. Stamatakis, Y. Mao, G. Northoff, Higher-order sensorimotor circuit of the brain’s global network supports human consciousness. Neuroimage. 231, 117850 (2021).

8. C. W. Wong, V. Olafsson, O. Tal, T. T. Liu, The amplitude of the resting-state fMRI global signal is related to EEG vigilance measures. Neuroimage. 83, 983–990 (2013).

9. C. W. Wong, P. N. DeYoung, T. T. Liu, Differences in the resting-state fMRI global signal amplitude between the eyes open and eyes closed states are related to changes in EEG vigilance. Neuroimage. 124, 24–31 (2016).

10. C. Chang, D. A. Leopold, M. L. Schölvinck, H. Mandelkow, D. Picchioni, X. Liu, F. Q. Ye, J. N. Turchi, J. H. Duyn, Tracking brain arousal fluctuations with fMRI. Proc. Natl. Acad. Sci. U. S. A. 113, 4518–4523 (2016).

11. S. Tanabe, Z. Huang, J. Zhang, Y. Chen, S. Fogel, J. Doyon, J. Wu, J. Xu, J. Zhang, P. Qin, X. Wu, Y. Mao, G. A. Mashour, A. G. Hudetz, G. Northoff, Altered Global Brain Signal during Physiologic, Pharmacologic, and Pathologic States of Unconsciousness in Humans and Rats. Anesthesiology. 132, 1392–1406 (2020).

12. C. Orban, R. Kong, J. Li, M. W. L. Chee, B. T. T. Yeo, Time of day is associated with paradoxical reductions in global signal fluctuation and functional connectivity. PLoS Biol. 18, e3000602 (2020).

13. X. Liu, J. A. De Zwart, M. L. Schölvinck, C. Chang, F. Q. Ye, D. A. Leopold, J. H. Duyn, Subcortical evidence for a contribution of arousal to fMRI studies of brain activity. Nat. Commun. 9, 1–10 (2018).

14. M. J. Redinbaugh, J. M. Phillips, N. A. Kambi, S. Mohanta, S. Andryk, G. L. Dooley, M. Afrasiabi, A. Raz, Y. B. Saalmann, Thalamus Modulates Consciousness via Layer-Specific Control of Cortex. Neuron. 106, 66-75.e12 (2020).

15. J. Tasserie, L. Uhrig, J. D. Sitt, D. Manasova, M. Dupont, S. Dehaene, B. Jarraya, Deep brain stimulation of the thalamus restores signatures of consciousness in a nonhuman primate model. Sci. Adv. 8, 1–18 (2022).

16. N. D. Schiff, J. T. Giacino, K. Kalmar, J. D. Victor, K. Baker, M. Gerber, B. Fritz, B. Eisenberg, J. O’Connor, E. J. Kobylarz, S. Farris, A. Machado, C. McCagg, F. Plum, J. J. Fins, A. R. Rezai, Behavioural improvements with thalamic stimulation after severe traumatic brain injury. Nature. 448, 600–603 (2007).

17. S. W. Hughes, D. W. Cope, K. L. Blethyn, V. Crunelli, Cellular Mechanisms of the Slow (<1 Hz) Oscillation in Thalamocortical Neurons In Vitro. Neuron. 33, 947–958 (2002).

18. J. F. A. Poulet, L. M. J. Fernandez, S. Crochet, C. C. H. Petersen, Thalamic control of cortical states. Nat. Neurosci. 15, 370–372 (2012).

19. F. David, J. T. Schmiedt, H. L. Taylor, G. Orban, G. Di Giovanni, V. N. Uebele, J. J. Renger, R. C. Lambert, N. Leresche, V. Crunelli, Essential thalamic contribution to slow waves of natural sleep. J. Neurosci. 33, 19599–19610 (2013).

20. X. Kong, R. Kong, C. Orban, P. Wang, S. Zhang, K. Anderson, A. Holmes, J. D. Murray, G. Deco, M. van den Heuvel, B. T. T. Yeo, Sensory-motor cortices shape functional connectivity dynamics in the human brain. Nat. Commun. 12, 6373 (2021).

21. X. Liu, J. H. Duyn, Time-varying functional network information extracted from brief instances of spontaneous brain activity. Proc. Natl. Acad. Sci. U. S. A. 110, 4392–4397 (2013).

22. J. Zhang, Z. Huang, S. Tumati, G. Northoff, Rest-task modulation of fMRI-derived global signal topography is mediated by transient coactivation patterns. PLoS Biol. 18, 1–22 (2020).

23. C.-C. Huang, E. T. Rolls, J. Feng, C.-P. Lin, An extended Human Connectome Project multimodal parcellation atlas of the human cortex and subcortical areas. Brain Struct. Funct. 227, 763–778 (2022).

24. S. A. Prescott, S. Ratté, in Conn’s Translational Neuroscience, P. M. Conn, Ed. (Elsevier, 2017), pp. 517–539.

25. S. Warren, N. F. Capra, R. P. Yezierski, in Fundamental Neuroscience for Basic and Clinical Applications: Fifth Edition, D. E. Haines, G. A. Mihailoff, Eds. (Elsevier., Fifth Edit., 2017), pp. 243-257.e1.

26. D. Liu, Y. Dan, A Motor Theory of Sleep-Wake Control: Arousal-Action Circuit. Annu. Rev. Neurosci. 42, 27–46 (2019).

27. J. Turchi, C. Chang, F. Q. Ye, B. E. Russ, D. K. Yu, C. R. Cortes, I. E. Monosov, J. H. Duyn, D. A. Leopold, The Basal Forebrain Regulates Global Resting-State fMRI Fluctuations. Neuron. 97, 940-952.e4 (2018).

28. M. L. Schölvinck, A. Maier, F. Q. Ye, J. H. Duyn, D. A. Leopold, Neural basis of global resting-state fMRI activity. Proc. Natl. Acad. Sci. U. S. A. 107, 10238–10243 (2010).

29. J. D. Power, M. Plitt, T. O. Laumann, A. Martin, Sources and implications of whole-brain fMRI signals in humans. Neuroimage. 146, 609–625 (2017).

30. M. D. Fox, A. Z. Snyder, J. L. Vincent, M. Corbetta, D. C. Van Essen, M. E. Raichle, The human brain is intrinsically organized into dynamic, anticorrelated functional networks. Proc. Natl. Acad. Sci. U. S. A. 102, 9673–9678 (2005).

31. M. Lee, L. R. D. Sanz, A. Barra, A. Wolff, J. O. Nieminen, M. Boly, M. Rosanova, S. Casarotto, O. Bodart, J. Annen, A. Thibaut, R. Panda, V. Bonhomme, M. Massimini, G. Tononi, S. Laureys, O. Gosseries, S.-W. Lee, Quantifying arousal and awareness in altered states of consciousness using interpretable deep learning. Nat. Commun. 13, 1064 (2022).

32. T. C. Gent, M. Bandarabadi, C. G. Herrera, A. R. Adamantidis, Thalamic dual control of sleep and wakefulness. Nat. Neurosci. 21, 974–984 (2018).

33. S. Ren, Y. Wang, F. Yue, X. Cheng, R. Dang, Q. Qiao, X. Sun, X. Li, Q. Jiang, J. Yao, H. Qin, G. Wang, X. Liao, D. Gao, J. Xia, J. Zhang, B. Hu, J. Yan, Y. Wang, M. Xu, Y. Han, X. Tang, X. Chen, C. He, Z. Hu, The paraventricular thalamus is a critical thalamic area for wakefulness. Science. 362, 429–434 (2018).

34. M. Falahpour, C. Chang, C. W. Wong, T. T. Liu, Template-based prediction of vigilance fluctuations in resting-state fMRI. Neuroimage. 174, 317–327 (2018).

35. G. R. Poudel, C. R. H. Innes, P. J. Bones, R. Watts, R. D. Jones, Losing the struggle to stay awake: Divergent thalamic and cortical activity during microsleeps. Hum. Brain Mapp. 35, 257–269 (2014).

36. J. L. Ong, D. Kong, T. T. Y. Chia, J. Tandi, B. T. Thomas Yeo, M. W. L. Chee, Coactivated yet disconnected-Neural correlates of eye closures when trying to stay awake. Neuroimage. 118, 553–562 (2015).

37. V. J. Kiviniemi, H. Haanpää, J. H. Kantola, J. Jauhiainen, V. Vainionpää, S. Alahuhta, O. Tervonen, Midazolam sedation increases fluctuation and synchrony of the resting brain BOLD signal. Magn. Reson. Imaging. 23, 531–537 (2005).

38. R. Huber, M. F. Ghilardi, M. Massimini, G. Tononi, Local sleep and learning. Nature. 430, 78–81 (2004).

39. V. V. Vyazovskiy, U. Olcese, Y. M. Lazimy, U. Faraguna, S. K. Esser, J. C. Williams, C. Cirelli, G. Tononi, Cortical Firing and Sleep Homeostasis. Neuron. 63, 865–878 (2009).

40. S. Vahdat, S. Fogel, H. Benali, J. Doyon, Network-wide reorganization of procedural memory during NREM sleep revealed by fMRI. Elife. 6, 1–24 (2017).

41. Z. Fang, L. B. Ray, A. M. Owen, S. M. Fogel, Brain activation time-locked to sleep spindles associated with human cognitive abilities. Front. Neurosci. 13, 1–16 (2019).

42. Z. Huang, X. Liu, G. A. Mashour, A. G. Hudetz, Timescales of intrinsic BOLD signal dynamics and functional connectivity in pharmacologic and neuropathologic states of unconsciousness. J. Neurosci. 38, 2304–2317 (2018).

43. J. T. Giacino, K. Kalmar, J. Whyte, The JFK Coma Recovery Scale-Revised: Measurement characteristics and diagnostic utility. Arch. Phys. Med. Rehabil. 85, 2020–2029 (2004).

