## Supplementary_Materials for "The Neural Correlates of Arousal: The Ventral Posterolateral Nucleus-Global Transient Co-Activation"

**This PDF file includes:**

Materials and Methods
Figs. S1 to S5

### Materials and Methods

Participants and Data Acquisition

Sleep Dataset

Two datasets were combined to form a large data. The first dataset was acquired at Institut Universitaire de Gériatrie de Montréal (IUGM) and the other was at Western University. All participants who passed inclusion/exclusion criteria were instructed to sleep in the scanner over night with a simultaneous EEG-fMRI recording. After excluding the participants who had not enough continuous segments (90 fMRI volumes) in each sleep stage, we collected a total of 30 participants in wakefulness (12 male, age 25.06 ± 4.08), 37 participants in N2 sleep (15 male, age 25.30 ± 4.18), and 18 participants in N3 sleep (8 male, age 24.06 ± 4.07). Details about the participants and data acquisition of the two datasets were as follows.

Sleep Dataset1: IUGM

Participants and experimental design. This dataset served as the basis for a published study (*40*). Thirteen healthy right-handed adults (5 male, age 27.4 ± 3.6) passed the inclusion/exclusion criteria, and a simultaneous EEG-fMRI recording scan took place while subjects slept in the scanner. The sleep session was terminated when the maximum possible number of volumes for a single fMRI session (4000 volumes, lasting a maximum of 2.5 hr) was reached, or if subjects voluntarily terminated the session. Ethical and scientific approval was obtained from the Research Ethics Board at the IUGM, Montreal, Quebec, Canada and an informed written consent was obtained prior to entering the study. See the previous study for more detailed information about the participants criteria (*40*).

Data acquisition. Images were collected using a 3T TIM TRIO Siemens scanner with a 12-channel head coil. A structural volume was acquired in the sagittal plane using a magnetization prepared rapid gradient echo (MPRAGE) sequence: TR = 2300 ms, TE = 2.98 ms, FA = 9˚, 176 slices, FoV = 256 × 256 mm2, voxel size = 1 × 1 × 1 mm3. For functional acquisitions, an echo-planar imaging (EPI) gradient echo sequence was used with the following parameters: TR = 2160 ms, TE = 30 ms, FA = 90˚, FoV = 220 × 220 mm2, matrix size = 64 × 64, 40 transverse slices, slice thickness = 3 mm, 10% inter-slice gap, inplane resolution = 3.44 × 3.44 mm2. In order to minimize the effects of gradient artifact on electroencephalography recordings, the sequence parameters were chosen so that the MR scan repetition time (2160 ms) matched a common multiple of the EEG sample time (0.2 ms), the product of the scanner clock precision (0.1 ms) and the number of slices (40 slices). NREM periods and intermittent wakefulness periods were classified by the EEG scoring according to standard criteria of the American Academy of Sleep Medicine (AASM), and the longest continuous (at least 90 fMRI volumes) segments of sleep stages were selected to avoid discontinuities in our data analysis. After discarding subjects with insufficient data length of NREM or wakefulness periods, there remained 11 subjects for wakefulness, 12 subjects for N2 sleep, and 8 subjects for N3 sleep. See the previous study for more detailed information about the simultaneous EEG-fMRI data acquisition and EEG data preprocesssing (*40*).

Sleep Dataset2: Western University

Participants and experimental design. This dataset served as the basis for a published study (*41*). Twenty-seven healthy right-handed adults (10 male, age 23.76 ± 3.72) passed the inclusion/exclusion criteria and slept in the scanner with a simultaneous EEG-fMRI recording. For each participant, up to 2 h of sleep EEG–fMRI data was acquired. All study procedures and methods adhered to the Declaration of Helsinki and were approved by the Western University Health Science research ethics board. See the previous study for more detailed information about criteria (*41*).

Data acquisition. Functional magnetic resonance imaging was performed at a 3.0T Magnetom Prisma MR imaging system (Siemens, Erlangen, Germany) using a 64-channel head coil. High-resolution anatomic images were acquired using a standard 3D Multislice MPRAGE sequence (TR = 2300 ms, TE = 2.98 ms, TI = 900 ms, FA = 9◦, 176 slices, FoV = 256 × 256 mm2, matrix size = 256 × 256 × 176, voxel size = 1 × 1 × 1 mm3). During the sleep session, T2 ∗-weighted fMRI images were acquired with a gradient echo-planar imaging (EPI) sequence using axial slice orientation (TR = 2160 ms, TE = 30 ms, FA = 90◦, 40 transverse slices, 3 mm slice thickness, 10% inter-slice gap, FoV = 220 × 220 mm2, matrix size = 64 × 64 × 40, voxel size = 3.44 × 3.44 × 3 mm3). An expert who was a registered polysomnographic technologist scored the EEG data acquired during the simultaneous EEG–fMRI sleep recordings according to standard criteria of the American Academy of Sleep Medicine (AASM). The longest continuous (at least 90 fMRI volumes) segments of sleep stages were selected to avoid discontinuities in our data analysis. After discarding subjects with insufficient data length of NREM or wakefulness periods, there remained 19 subjects for wakefulness, 25 subjects for N2 sleep, and 10 subjects for N3 sleep. See the previous study for more detailed information about the simultaneous EEG-fMRI data acquisition and EEG data preprocesssing (*41*).

Anesthesia dataset

Participants and experimental design. This dataset served as the basis for a published study (*42*). A total of 15 healthy adults (9 male, age 26.73 ± 4.79) were included in this study. All subjects received four 15-min resting-state scans in wakefulness, propofol-induced light and deep sedation, and recovery. Two levels of responsiveness were targeted: light sedation, in which volunteers showed lethargic response to verbal commands (observer's assessment of alertness/sedation, OAAS, score 4), and deep sedation, during which volunteers showed no response to verbal commands (OAAS score 2-1). The corresponding target plasma concentrations vary across subjects (light sedation: 0.98 ± 0.18 μg/ml; deep sedation: 1.88 ± 0.24 μg/ml) because of the variability in individual sensitivity to anesthetics. The Institutional Review Board of Medical College of Wisconsin approved the experimental protocol. In this study, we combined the first half of wakefulness scan (first 7.5 min) and the second half of recovery scan (second 7.5 min) to generate a full conscious scan, which avoided the possible asleep state in a long resting state, and the unconscious state at the beginning of recovery. And we adopted the combined wakefulness and deep sedation scans in the current study. After excluding the subjects who had exceeding head motion, 14 subjects in wakefulness (8 male, age 26.57 ± 4.93) and 12 subjects in deep sedation (6 male, age 25.91 ± 4.70) remained.

Data acquisition. Image data were acquired using a 3T Signa GE 750 scanner (GE Healthcare) with a standard 32-channel transmit/receive head coil. Functional imaging data were acquired using gradient-echo EPI images of the whole brain: 41 slices, TR/TE = 2000/25 ms, slice thickness = 3.5 mm, in-plane resolution = 3.5 × 3.5 mm; FOV = 224 mm, flip angle = 77°, image matrix: 64 × 64. High-resolution spoiled gradient-recalled echo anatomical images were acquired before the functional scans with parameters: TE/TR/TI, 8.2/3.2/450 ms, slice thickness = 1 mm, number of slices = 150, flip angle = 12°, field of view = 24 cm, matrix size = 256 × 256. More detail information about the experiment protocol and data acquisition parameters were shown in the previous study (*42*).

EO/EC dataset

This dataset combined two open resting-state datasets, which were recruited from Beijing Normal University. All participants were instructed to rest with eyes open or eyes closed without falling asleep. After excluding the participants with incomplete image data, we obtained a total of 67 participants (34 male, age 22.04 ± 2.18) in this study. More information about the participants and data acquisition of these datasets were introduced in the following.

Dataset1: Beijing Normal University 1

Participants and experimental design. This dataset is recruited in Beijing Normal University, which is available on the NITRC (NeuroImaging Tools and Resources Collaboratory) website (http://fcon_1000.projects.nitrc.org/indi/retro/BeijingEOEC.html). A total of 47 healthy participants (23 male, age 22.51 ± 2.17) were included after excluding one participant with incomplete data. None of these participants had a history of medical, neurological, or psychiatric disorders. Each participant received three resting-state fMRI scans and was instructed to keep as motionless as possible and not to engage in any systematic thinking. Each of these scans lasted for 8 minutes. During the first scan, each participant was instructed to rest with their eyes closed. The second and third resting-state scans consisted runs with both eyes open or closed, of which the order was counterbalanced across all participants. Immediately after each session, the experiment operator spoke briefly with the participants. All the participants reported that they had not fallen asleep during the scan. Only the second and third resting scans were used in this study.

Data acquisition. The fMRI data were acquired on a 3T Siemens Trio TIM MR scanner using a gradient-echo EPI sequence: TR = 2000 ms, TE = 30 ms, flip angle = 90°, FOV = 200 × 200 mm2; matrix = 64 × 64; slice thickness/gap = 3.5/0.7 mm; 33 slices. A total of 240 volumes were acquired in each run (8 min). A high-resolution T1-weighted anatomical image was also acquired using the MP-RAGE sequence: 128 slices, TR = 2530 ms, TE = 3.39 ms, slice thickness/gap = 1.33/0 mm, flip angle = 7º, inversion time = 1100 ms, FOV = 256 × 256 mm2, and in-plane resolution = 256 × 192.

Dataset2: Beijing Normal University 2

Participants and experimental design. This dataset was recruited from Beijing Normal University, which is available on the NITRC (NeuroImaging Tools and Resources Collaboratory) website (http://rfmri.org/BeijingEOEC2_Raw). A total of 20 healthy participants (10 male, age 20.95 ± 1.82) were included. None of these participants had a history of medical, neurological, or psychiatric disorders. The participants received four scans, including three resting-state and one visual response task. They first underwent an EC resting-state scan. The second and third resting-state scan were in EO and EO-F (EO with a fixation), respectively. These scans were counterbalanced across the participants. Each of these scans lasted for 8 minutes. During the three resting-state sessions, the participants were instructed to keep as motionless as possible and not to engage in any systematic thinking. Immediately after each scanning session, the experiment operator had a short communication with the participants. All participants reported that they had not fallen asleep during the scan. Only the scans of EC and EO were used in this study.

Data acquisition. The fMRI data were acquired on a 3T Siemens Trio TIM MR scanner using an echo-planar imaging sequence with the following parameters: TR = 2000 ms; TE = 30 ms; flip angle = 90°; FOV = 200 × 200 mm; in-plane resolution = 64 × 64; slice thickness/gap = 3/0.6 mm; 33 slices. A total of 240 volumes were acquired in each run (8 min). In addition, a T1-weighted sagittal three-dimensional magnetization-prepared rapid gradient echo (MPRAGE) sequence was acquired, covering the entire brain: 128 slices, TR = 2530 ms, TE = 3.39 ms, slice thickness = 1.33 mm, flip angle = 7º, inversion time = 1100 ms, FOV = 256 × 256 mm2, and in-plane resolution = 256 × 192.

UWS Dataset

The Zhujiang Hospital Dataset

Participants and experimental design. This dataset included 23 UWS patients with well-preserved brain structures and 21 healthy subjects. The UWS patients were assessed using the Coma Recovery Scale-Revised (CRS-R) (*43*) before the fMRI scanning. Two hundred and forty fMRI volumes (~8 min) were acquired during rest for each subject. All participants were given the same instructions, in which they were told to take a comfortable supine position, relax, and not concentrate on anything in particular during the scanning. Informed written consent was obtained from the patients' legal representatives. The study was approved by the Ethics Committee of Zhujiang Hospital, Guangzhou, China. After excluding the patients who had exceeding motion, 21 healthy subjects (10 male, age 32.33 ± 10.42 ) and 20 UWS patients ( 10 male, age 52.55 ± 11.64) remained.

Data acquisition. Image data were acquired on the same Philips 3 Tesla scanner. Functional images were acquired using a T2*-weighted EPI sequence: TR/TE = 2000 ms/30 ms, FOV = 224 × 224 mm2, matrix = 64 × 64, 33 slices thickness = 3.5 mm, gap = 0.7 mm. A high-resolution T1-weighted anatomical image was also acquired for all participants with parameters: TE/TR = 3.2/7 ms, slice thickness = 1 mm, matrix size = 256 × 256.

The Shanghai Hospital Dataset

Participants and experimental design. This dataset served as the basis for a published study (*7*). We included 50 patients with structurally well-preserved brains under three conditions: UWS, MCS and BI with full conscious and a brain injury history. These structurally well-preserved patients, who were selected by author XW and then checked by author PQ according to their structural images, had limited brain lesions and limited brain structure distortion. The UWS and MCS patients were assessed using the Coma Recovery Scale-Revised (CRS-R) (*43*) before the fMRI scanning. Two hundred fMRI volumes (~6.7 min) were acquired during rest for each subject. All participants were given the same instructions, in which they were told to take a comfortable supine position, relax, close their eyes, and not concentrate on anything in particular during the scanning. Informed written consent was obtained from the patients' legal representatives. The study was approved by the Ethics Committee of Shanghai Huashan Hospital, Fudan University, Shanghai, China. In this study, only the UWS and BI patients were included.

After excluding the patients who had exceeding motions, 15 BI patients (11 male, age 38.8 ± 16.89) and 19 UWS patients (14 male, age 45.84 ± 12.49) remained.

Data acquisition. Image data were acquired on the same Siemens 3 Tesla scanner. Functional images were acquired using a T2*-weighted EPI sequence: TR/TE/θ = 2000 ms/35 ms/90°, FOV = 256 × 256 mm, matrix = 64 × 64, 33 slices thickness = 4 mm, gap = 0 mm. A high-resolution T1-weighted anatomical image was also acquired for all participants with parameters: TE/TR = 2.98/2300 ms, slice thickness = 1 mm, matrix size = 256 × 256. More detailed information about the experiment protocol and data acquisition parameters were shown in the previous study (*7*).

### Method

#### Data Preprocessing

All MRI images of each dataset were processed using the AFNI software package (https://afni.nimh.nih.gov/). First, the first three volumes were excluded, followed by the elimination of spikes and the correction of slice timing. All functional images were registered to the base volume, which was considered to have minimum outliers. Both the anatomical and functional images were then aligned and registered to standard MNI space, and the spatial resolution of each voxel was resampled into $3mm^3$. Subsequently, the functional images were smoothed using a 6-mm full-width at half-maximum (FWHM) Gaussian kernel. The functional data were then band-pass filtered (0.01–0.1 Hz), and the nuisance factors (six demeaned motion parameters, first differences of six motion parameter, and five principal components of the signals from white matter and cerebrospinal fluid) were regressed out. Moreover, motion was quantified as the Euclidean norm calculated from the six motion parameters for 2 consecutive TRs, and a displacement of > 0.4 mm was considered excessive. The volume at the corresponding time point of the excessive motion, as well as the preceding volume, were replaced by interpolated neighboring non-excessive volumes. Furthermore, volumes with >10% voxels of the base volume were flagged as outliers were also replaced with the interpolation method. Finally, the time series of each voxel was z-score transformed by subtracting its temporal mean and dividing by its temporal standard deviation.

#### GS Co-activation

We used the FSL software (https://fsl.fmrib.ox.ac.uk/fsl/fslwiki) to segment the anatomical image and obtained gray matter (GM) for each participant. The GS was extracted by averaging the z-score transformed signals within GM. The time points of the GS in the top 17% were selected to represent the instantaneous increase (*13*, *22*). To control the effect of various arousal states on the whole brain, and to highlight the relative relation between GS and other regions, the z-score map of each brain volume was normalized by being subtracted and divided by the GS value at that volume. Finally, the normalized activation maps at the time points of GS increase (top 17%) were averaged to generate the GS co-activation map (*13, 22*). (Fig.1B).

To compare the voxel-wise difference of GS co-activation maps between conditions, we used ANOVA for the sleep dataset, unpaired t-test for the anesthesia dataset (wakefulness vs. deep sedation), and paired t-test the for EO/EC dataset (EO vs. EC). The voxel-wise significance of all these comparisons were $p<0.001$ and were corrected in a cluster level for multicomparisons with alpha < 0.005, using 3dClustSim in AFNI. The regions with a significant difference were then chosen for subsequent ROI analysis.

For the sleep dataset, unpaired t-test of ROIs was conducted in each paired comparison (W vs. N1, W vs. N2, N2 vs. N3). For the anesthesia dataset, unpaired t-test was also used for comparison between conditions. For the EO/EC dataset, we used paired t-test for conditions comparison. For the DOC datasets, we compared the conditions with unpaired t-test on the overlapped thalamus region found in the sleep, anesthesia and EO/EC datasets. And the multi-comparison correction were conducted using Bonferroni method.

#### Normalized Activation at Random Time Points

To test whether the ROIs showed a distinctive activation pattern at GS increases, we examined their normalized activation at random time points. The number of the random time points was the same as in the GS increases (top 17%). The normalized values of activation in the ROIs were then averaged across random time points and were compared between conditions in each dataset. All the group statistics were the same as previous ROI analysis in GS co-activation. And the multi-comparison correction were conducted using Bonferroni method.

#### GS co-activation of Physiological Noise

In order to exclude the contribution of physiological noise, such as WM and CSF, to the GS co-activation, we examined whether the GS co-activation of these noise were affected by altered arousal levels in each dataset. Specifically, for each participant, we adopted the masks of WM and CSF which is generated by setting the threshold of 95% on tissue probability map in SPM. And the GS co-activation were averaged within these masks. We then compared the GS-activation of WM and CSF between conditions in each dataset. All the group statistics were the same as previous ROI analysis in GS co-activation. And the multi-comparison correction were conducted using Bonferroni method.

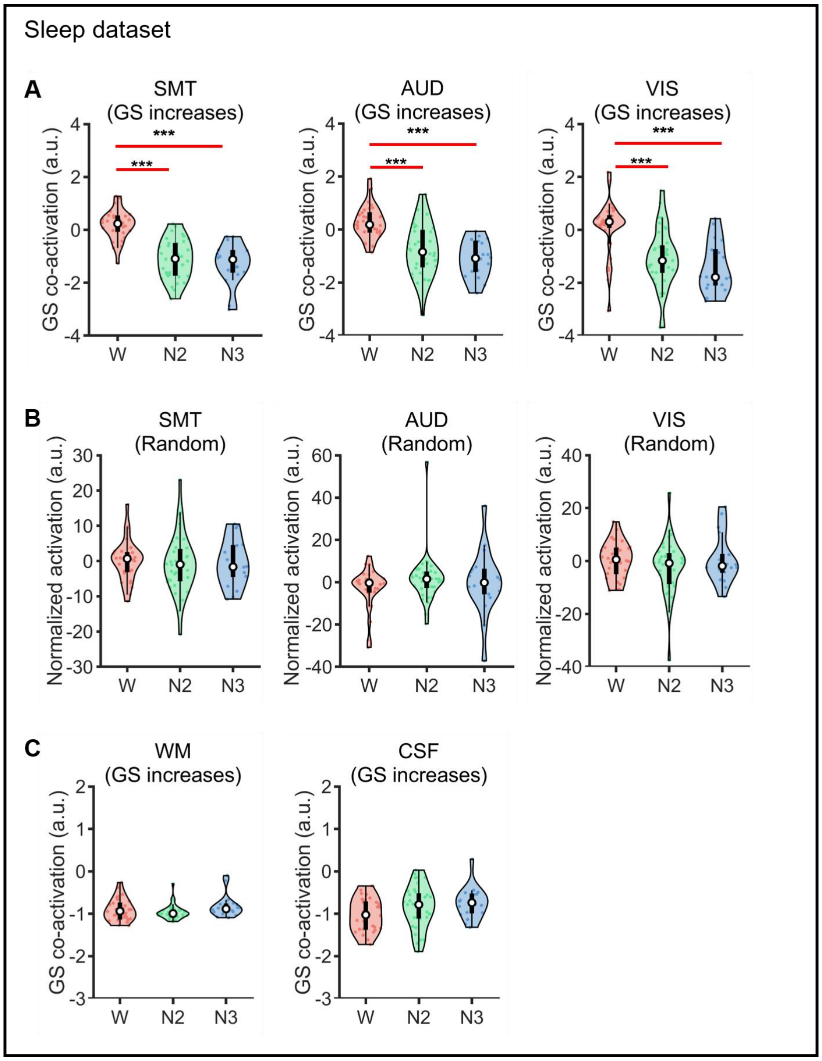


**Fig. S1.** (**A**) Comparison of GS co-activation between sleep stages at SMT, AUD and VIS. (**B**) Comparison of normalized activation during random time points between sleep stages at SMT, AUD and VIS. (**C**) Comparison of GS co-activation between sleep stages at WM and CSF

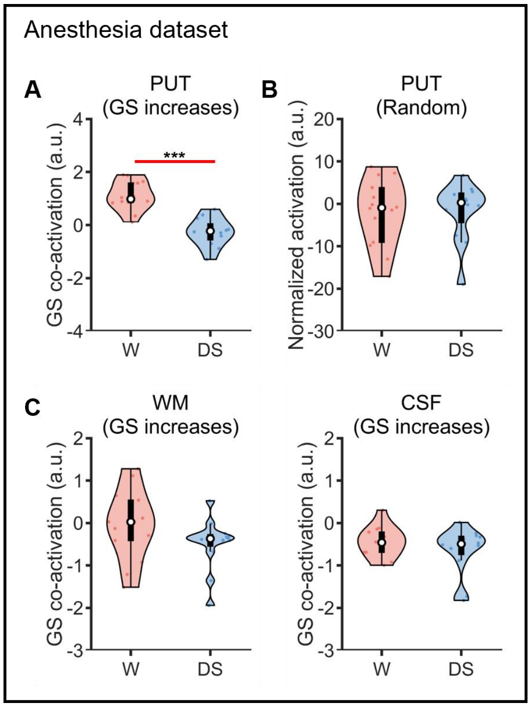


**Fig. S2.** (**A**) Comparison of GS co-activation between W and DS at putamen (PUT). (**B**) Comparison of normalized activation during random time points between W and DS at PUT. (**C**) Comparison of GS co-activation between W and DS at WM and CSF.

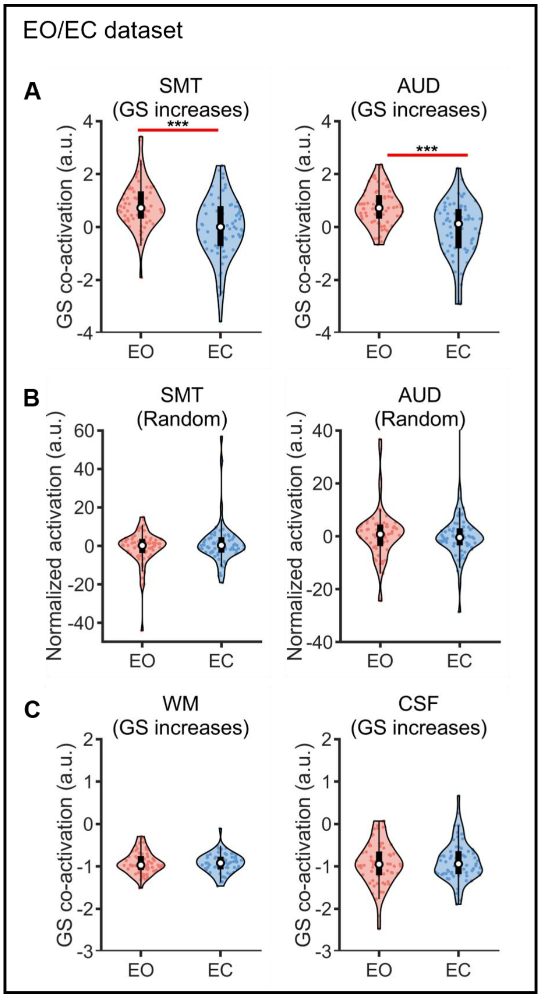


**Fig. S3.** (**A**) Comparison of GS co-activation between EO and EC at SMT and AUD. (**B**) Comparison of normalized activation during random time points between EO and EC at SMT and AUD. (**C**) Comparison of GS co-activation between EO and EC at WM and CSF.

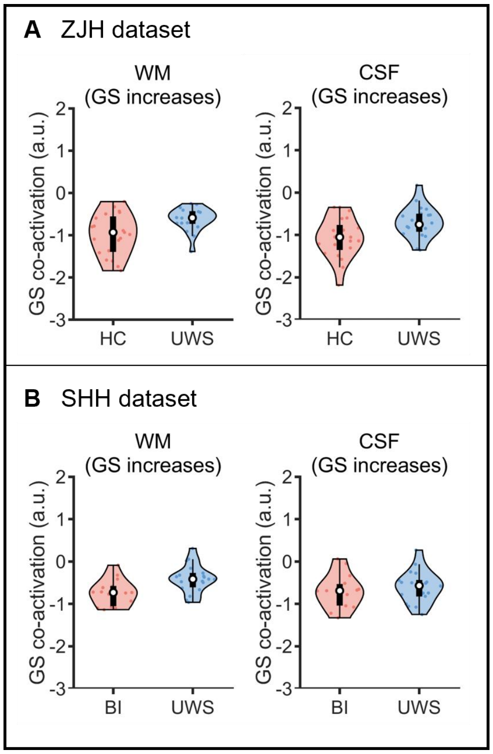


**Fig. S4.** (**A**) ZJH dataset: comparison of GS co-activation between HC and UWS at WM and CSF. (**B**) SHH dataset: comparison of GS co-activation between BI and UWS at WM and CSF.

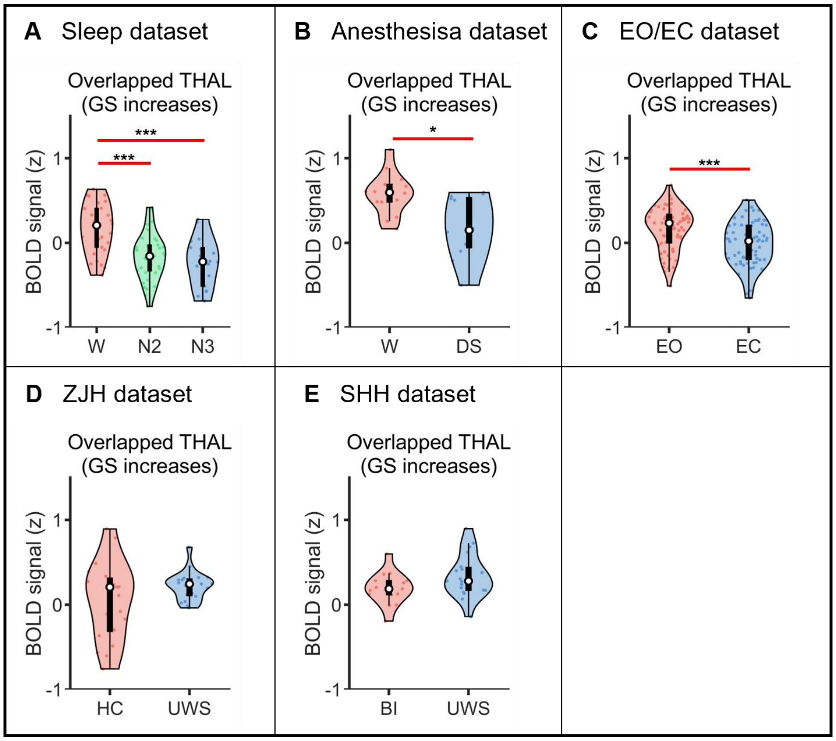


**Fig. S5.** Comparison of the averaged BOLD signal during GS increases at the overlapped thalamus regions in all datasets.
